# An invariant *Trypanosoma vivax* vaccine antigen inducing protective immunity

**DOI:** 10.1101/2021.02.10.430711

**Authors:** Delphine Autheman, Cécile Crosnier, Simon Clare, David A. Goulding, Cordelia Brandt, Katherine Harcourt, Charlotte Tolley, Francis Galaway, Malhar Khushu, Han Ong, Alessandra Romero Ramirez, Craig W. Duffy, Andrew P. Jackson, Gavin J. Wright

**Affiliations:** Cell Surface Signalling Laboratory, Wellcome Sanger Institute, Hinxton, Cambridge, CB10 1SA UK; Pathogen Support Team, Wellcome Sanger Institute, Hinxton, Cambridge, CB10 1SA UK; Electron and Advanced Light Microscopy, Wellcome Sanger Institute, Hinxton, Cambridge, CB10 1SA UK; Department of Infection Biology, University of Liverpool, 146 Brownlow Hill, Liverpool, L3 5RF UK; Department of Biology, Hull York Medical School, York Biomedical Research Institute, University of York, Wentworth Way, York, YO10 5DD, UK

## Abstract

Trypanosomes are protozoan parasites that cause infectious diseases including human African trypanosomiasis (sleeping sickness), and nagana in economically-important livestock animals^1,2^. An effective vaccine against trypanosomes would be an important control tool, but the parasite has evolved sophisticated immunoprotective mechanisms including antigenic variation^3^ that present an apparently insurmountable barrier to vaccination. Here we show using a systematic genome-led vaccinology approach^4^ and a murine model of *Trypanosoma vivax* infection^5^ that protective invariant subunit vaccine antigens can be identified. Vaccination with a single recombinant protein comprising the extracellular region of a conserved cell surface protein localised to the flagellum membrane termed “invariant flagellum antigen from *T. vivax*” (IFX) induced long-lasting protection. Immunity was passively transferred with immune serum, and recombinant monoclonal antibodies to IFX could induce sterile protection and revealed multiple mechanisms of antibody-mediated immunity, including a major role for complement. Our discovery identifies a vaccine candidate for an important parasitic disease that has constrained the socioeconomic development of sub-Saharan African countries^6^ and challenges long-held views that vaccinating against trypanosome infections cannot be achieved.

## Main

African trypanosomiasis is an infectious disease caused by unicellular parasites of the genus *Trypanosoma* that are transmitted by the bite of an infected tsetse fly. In humans, trypanosome infections cause sleeping sickness: a deadly disease that threatens the lives of millions of people living in over 30 sub-Saharan countries^7^. Some species of trypanosome also infect important livestock animals such as cattle, goats and pigs causing the wasting disease nagana which affects the livelihoods of people relying on these animals for milk, food and draught power^1^. Approximately three million cattle die from this disease every year^8^ resulting in an estimated direct annual economic impact of many hundreds of millions of dollars^9^. This disease therefore represents a major barrier for the socioeconomic advancement of many African countries which are dependent on smallholders to produce the vast majority of food. Nagana (animal African trypanosomiasis) is primarily caused by *T. vivax* and *T. congolense* and is currently managed with drugs, but these are not satisfactory due to their side effects, and drug resistance is increasing^10^. These parasites have a broad host range, including wild animals that creates a constant reservoir, making eradication impractical^11^. Vaccination against trypanosomes has long been considered unachievable due to the evolution of parasite immune evasion strategies that enable it to survive in host blood. These strategies include antigenic variation: the serial expression of an abundant allelically-excluded variable surface glycoprotein (VSG), and the rapid removal of surface-bound antibodies by hydrodynamic sorting^3,12^. It is widely thought that the VSG forms a constantly-changing impenetrable surface coating that sterically shields other surface proteins from host antibodies leading to chronic infections characterised by oscillating parasitaemia; however, a careful analysis of recent structural information suggests that this model may not fully explain the protective role of the VSGs^13^. We therefore hypothesized that the VSGs may also subvert natural immunity by preventing the acquisition of high titre antibody responses to protective antigens suggesting that eliciting prophylactic unnatural host immunity by vaccination could be achieved. We now report the identification of a conserved cell surface protein that we term “invariant flagellum antigen from *T. vivax*” (IFX) that when used as a subunit vaccine in a murine model of *T. vivax* infection, is capable of eliciting highly protective immunity.

## Results

### IFX induces immunity to *T. vivax*

To identify subunit vaccine candidates for *T. vivax,* we first established a genome-led reverse vaccinology approach^4^ using a murine infection model (Fig. 1a). This took advantage of a luciferase-expressing *T. vivax* parasite cell line selected at the Institut Pasteur^5,14^. By comparing the infection parameters of different doses and infection routes, we determined that adoptive transfer of ~100 parasites intravenously into BALB/c hosts resulted in a highly reproducible acute model of infection that permitted sensitive and accurate parasitaemia quantification using bioluminescent imaging as described^15^. We compiled a list of potential subunit vaccine candidates by searching the *T. vivax* genome^16^ for genes encoding predicted cell surface and secreted proteins that are likely to be accessible to vaccine-elicited antibodies. We preferentially selected candidates using the following criteria: 1) they did not belong to paralogous gene families to minimise the risk of functional redundancy in different mammalian hosts^17^; 2) contained >300 amino acids in their predicted extracellular region and so are likely to be accessible at the parasite cell surface; and, 3) had evidence of expression in the blood stages^18,19^. Using these criteria, 60 candidates were selected and numbered (Supplementary Table 1). Gene sequences encoding the entire predicted extracellular region of selected candidates were synthesised and cloned into a mammalian protein expression plasmid containing an exogenous secretion peptide and purification tags. Candidates were expressed as soluble recombinant proteins in mammalian HEK293 cells to increase the chances that structurally critical posttranslational modifications were added and therefore elicit host antibodies that recognize native antigens displayed by the parasite. Of the 60 expression plasmids tested, 39 yielded sufficient protein after purification for vaccination trials (Extended Data Fig. 1a). For vaccination, we selected a prime and two boost regime using alum as an adjuvant to bias host responses towards humoral immunity. To reduce any systemic adjuvant-elicited effects on disease progression, vaccinated animals were rested for a minimum of four weeks following the final boost before parasite challenge (Extended Data Fig. 1b, c). In preliminary experiments, we observed that *T. vivax* lost virulence once removed from donor animals, and so to avoid confounding effects due to the loss of parasite viability during the infection procedure, we ensured that infections were comparable in control animals challenged before and after the vaccinated animals (Extended Data Fig. 1b).

**Figure 1.**
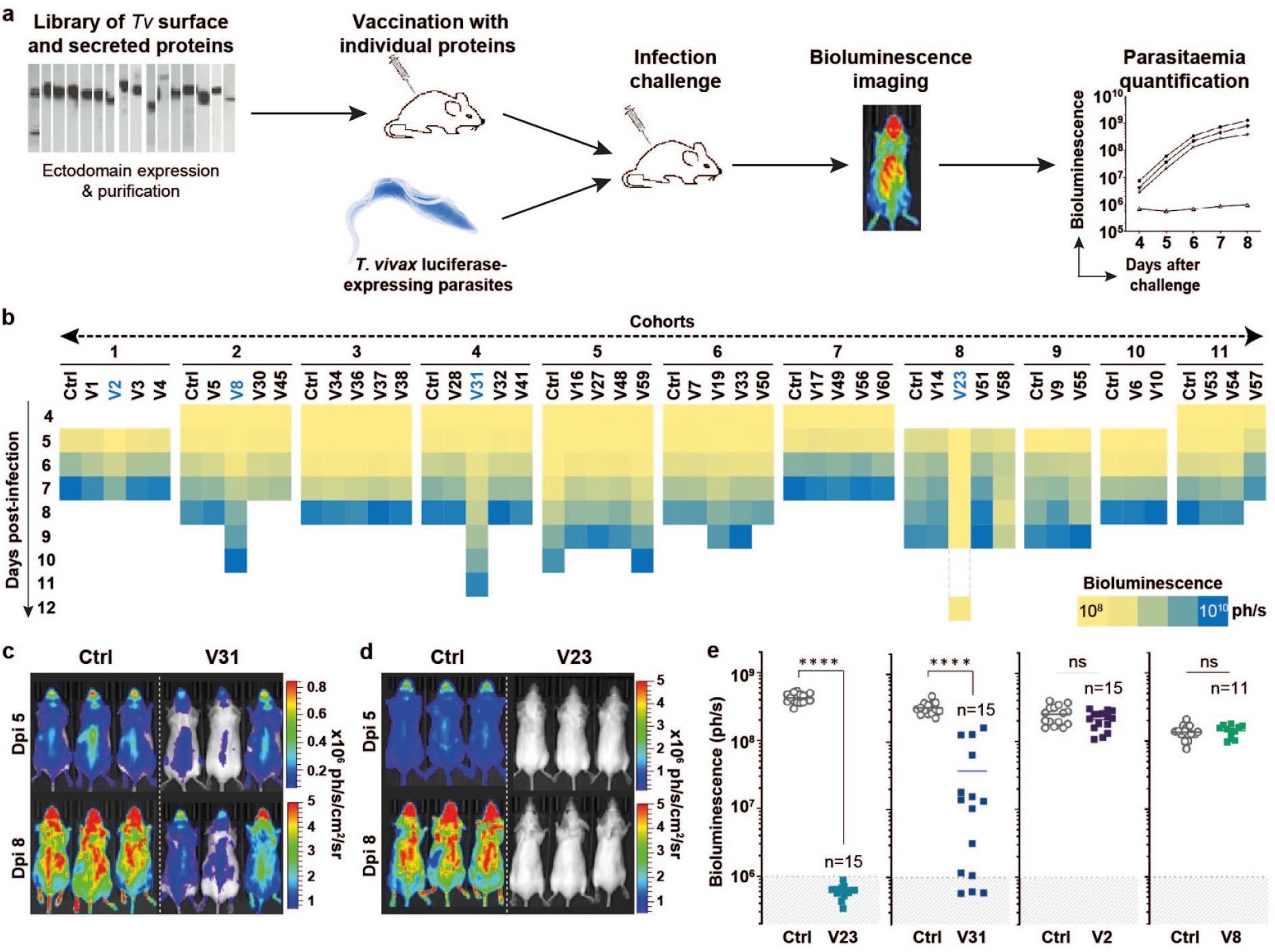
IFX/V23 induces protective immunity in a *Trypanosoma vivax* infection model. **a,** Schematic illustrating the genome-led vaccinology approach. **b,** Summary of mean parasitaemia (n ≥ 3 animals) quantified by bioluminescence in cohorts of vaccinated mice challenged with *T.* vi*vax*. **c**, **d**, Bioluminescent images of adjuvant-only control and mice vaccinated with V23 and V31 five and eight days post infection (Dpi). **e**, Quantification of replicate vaccinations and *T. vivax* infections with four antigens showing protective effects in the initial screen. Parasitaemia was quantified on day six using bioluminescence; data points represent individual animals and bars indicate mean ± SD. ns = not significant, **** P ≤ 0.00001 two-tailed t-test; grey shading indicates background bioluminescence thresholds.

Elicited antibody titres to each antigen were determined after the final boost and, as expected with a panel of different antigens, varied considerably with the vast majority (90%) having mean half-maximal responses at serum dilutions greater than 1:10,000 (Extended Data Fig. 1d). We found that of the 39 antigens tested, 34 had no effect on the infection parameters relative to controls (Fig. 1b, Extended Data Fig. 2). Statistically significant effects on parasite growth was observed with four antigens (Fig. 1b): two candidates (V2, V8) exhibited a slight, and one (V31) a longer delay to the ascending phase of parasitaemia (Fig. 1b, c), and one candidate (V23) showed no detectable parasites in all five vaccinated animals (Fig. 1b, d). One candidate, V58, although having lower mean parasitaemia relative to controls on day 8 rebounded on day 9 and so was not considered further (Fig. 1b, Extended Data Fig. 2). For each of the four candidates, experiments were repeated using independent protein preparations and larger cohorts of animals. The two candidates that induced a slight delay (V2, V8) did not replicate and so were not pursued further (Fig. 1e). Candidate V31 reduced the rate of parasite multiplication once more, inducing improved protection with 9/15 animals surviving until day 16 post infection (Fig. 1e). Again, V23 vaccination elicited robust protection (Fig. 1e), and longitudinal sampling of these animals showed that 10 out of the 15 animals were protected beyond at least day 170 (Extended Data Fig. 3). Dissection of protected animals several months after infection revealed no detectable extravascular reservoirs of parasites (Extended Data Fig. 4). Based on these and subsequent findings, we propose to name the V23 candidate (TvY486_0807240) IFX for “invariant flagellum antigen from *T. vivax*”.

### IFX localises to the flagellum

Sequence-based methods predict that IFX is a previously uncharacterised type I cell surface glycoprotein containing a short (18 amino acid) cytoplasmic region. IFX does not include any known protein domains and has no paralogs (protein sequence identity >25%) within *T. vivax* nor homologs in other sequenced *Trypanosoma* spp. genomes. *T. vivax* cannot be continuously cultured *in vitro* preventing gene targeting, and so to begin the functional characterisation of IFX, we asked whether it had a specific localisation in blood-stage parasites. Immunocytochemistry showed that staining was localised along the length of the flagellum and loosely concentrated in discrete puncta (Fig. 2a, Extended Data Fig. 5a). We used immunogold electron microscopy to determine the subcellular localisation in more detail. In transverse sections, IFX was enriched at the boundaries of where the flagellum is attached to the cell body; in different sections, these clusters were either uni- or bilaterally located (Fig. 2b). In mid-sagittal sections, IFX was located along the length of the flagellum membrane and concentrated in discrete clusters at the points where the flagellum was in close apposition to the cell membrane (Fig. 2c). Specifically, the gold particles were located between the flagellum and cell body membranes (Fig. 2d). We quantified these locations by counting the density of gold particle distribution in electron micrographs (Fig. 2e, Extended Data Fig. 5b). These data suggested IFX was localised to the flagellum membrane and particularly enriched as continuous or punctuated bilateral stripes along the flagellum, bordering the region where the flagellum is attached to the parasite cell body, suggesting a structural role in maintaining flagellar function^20^.

**Figure 2.**
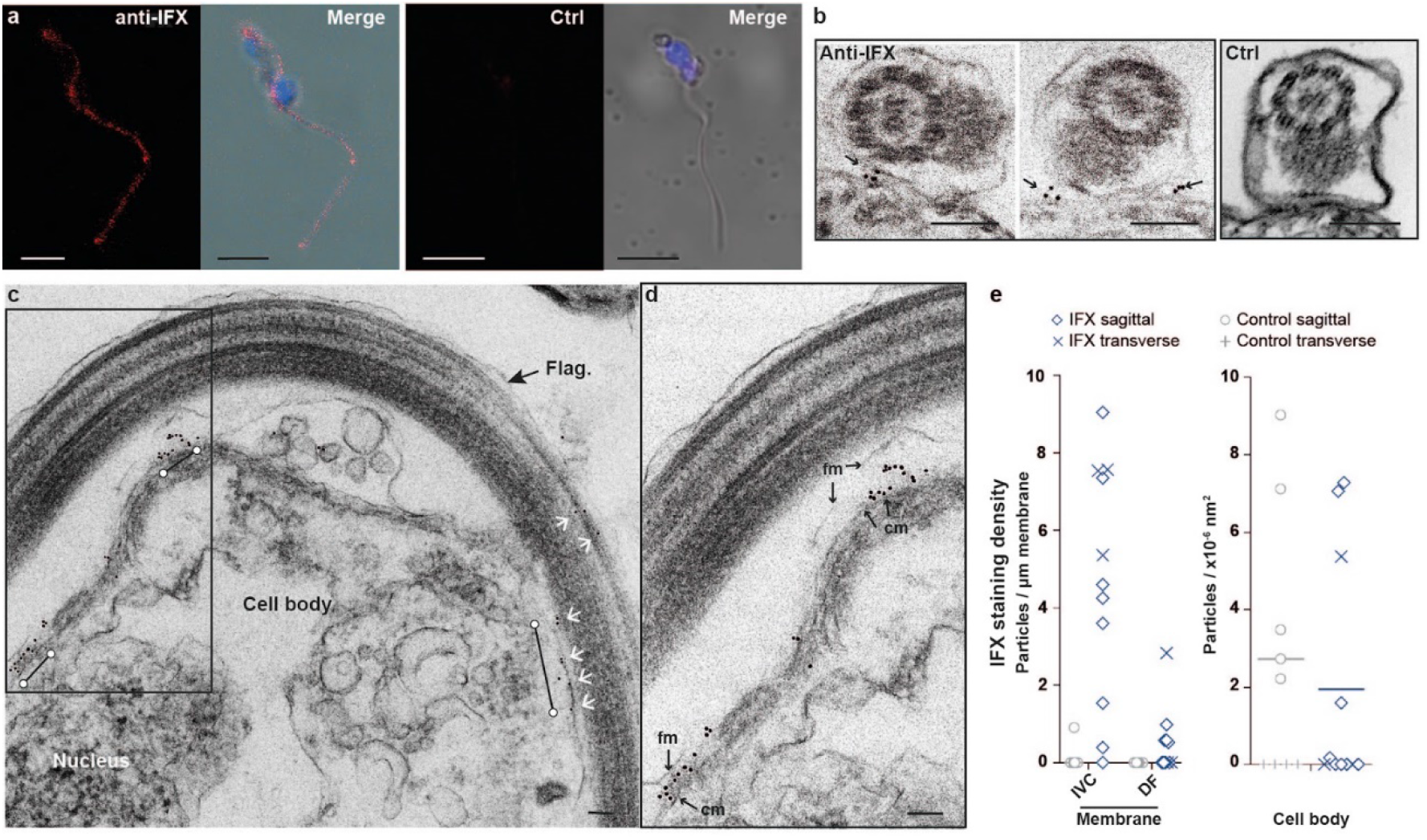
IFX is expressed on the *Trypanosoma vivax* flagellum membrane and concentrated at the periphery of flagellum-cell body contact. **a,** Immunofluorescence staining of *T. vivax* with rabbit anti-IFX antiserum (red, left panel) or control pre-immune serum (right), counterstained with DAPI (blue) demonstrates localisation of IFX to the flagellum. Scale bars = 5 μm. **b,** Immunogold electron microscopy using an anti-IFX mouse monoclonal localised IFX to the borders of where the flagellum is in contact with the parasite cell body in transverse sections (black arrows in two leftmost images) compared to isotype-matched control (right). **c,** IFX was concentrated in clusters along the length of the flagellum in mid-sagittal sections (white arrows and bars). **d**, Zoom of box in **c**, showing IFX located between the flagellum and cell membranes. Scale bars in (**b**), (**c**), and (**d**) = 100 nm. **e,** Anti-IFX particle staining density was quantified along the membrane interface of the ventral flagellum/cell body (IVC), dorsal flagellum (DF) and cell body area on sagittal and transverse sections. Bars represent means. flag. = flagellum; fm = flagellar membrane; cm = parasite cell membrane.

### Antibodies to IFX passively protect

To determine the immunological mechanisms of IFX-mediated protection, we first demonstrated that antibodies contributed to immunity by transferring immune serum from IFX-vaccinated animals to naïve recipients which inhibited parasite growth in a dose-dependent manner (Fig. 3a). Protection was not as potent compared to vaccinated animals, and this corresponded with lower titres of anti-IFX antibodies. Depletion of CD4 and CD8-positive T-lymphocytes and NK1.1-positive natural killer cells in IFX-vaccinated mice did not affect protective efficacy demonstrating these cell types were not direct executors of immunity once established (Extended Data Fig. 6). To further investigate the role of antibodies in immunity using an independent approach, we selected six hybridomas secreting monoclonal antibodies (mAbs) to IFX. Out of the six mAbs selected, three affected parasite growth when used in passive protection experiments (Fig. 3b). We determined the approximate location of the mAb binding sites on IFX and quantified their binding affinities but did not observe a simple positive correlation between their protective efficacy and either the location of their epitope or binding affinity (Extended Data Figure 7). The inhibitory effects of one selected antibody (8E12) titrated with dose (Fig. 3c).

**Figure 3.**
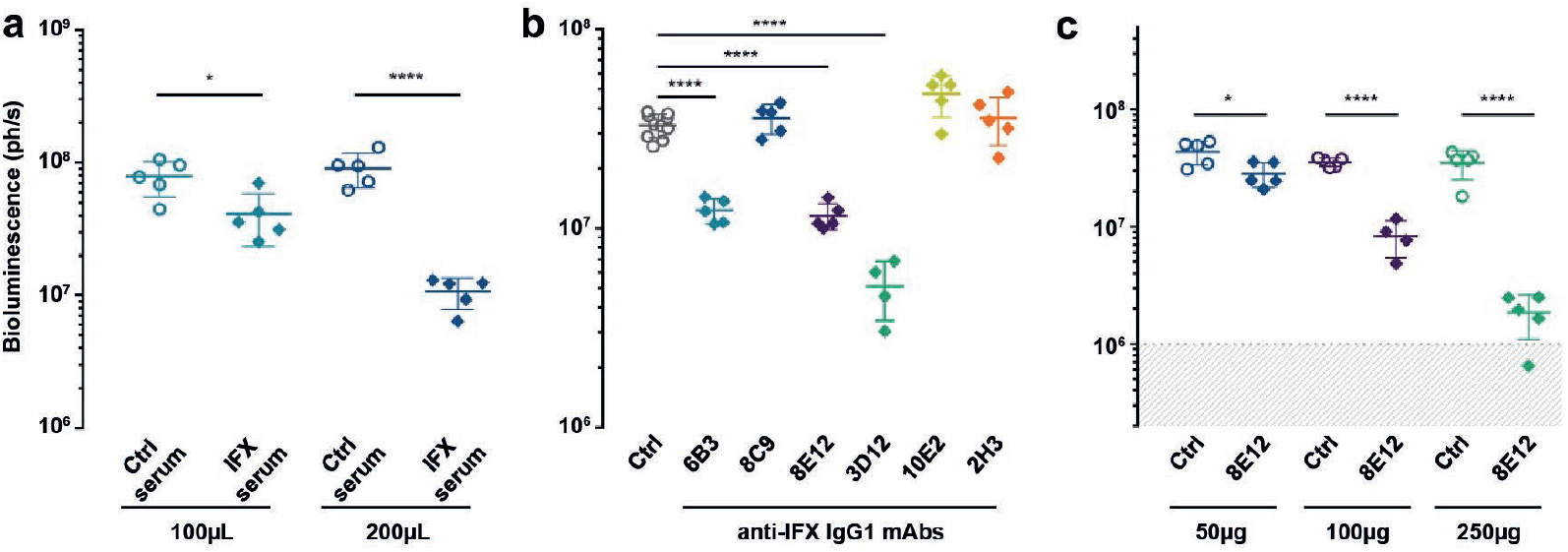
Passive transfer of immunity to *Trypanosoma vivax* infections with anti-IFX antibodies. **a,** Dose-dependent inhibition of *T. vivax* by adoptive transfer of sera from IFX-vaccinated mice relative to unimmunised control sera. **b**, Three of six anti-IFX IgG1-isotype mAbs each given at a dose of 3 x 100 μg passively protect against *T. vivax* infection relative to an isotype-matched control. **c**, Passive protection of the 8E12-IgG1 mAb is dose dependent. Parasitaemia was quantified at day 5 using bioluminescence and data points represent individual animals; bars indicate mean ± SD, groups were compared by one-way ANOVA with Sidak post-hoc test * P ≤ 0.01, **** P ≤ 0.00001; background bioluminescence threshold is indicated by grey shading.

### Multiple mechanisms of anti-IFX protection

Isotyping the mAbs to IFX revealed that they were all of the IgG1 subclass, which in mice, do not effectively recruit immune effector functions such as complement or bind activating Fc receptors with high affinity^21^ suggesting that direct antibody binding to IFX affected parasite viability. To establish the role of Fc-mediated immune effectors in IFX-mediated antibody protection, we selected a mAb, 8E12, that gave intermediate protective effects, and by cloning the rearranged antibody variable regions, switched the mAb isotype from IgG1 to IgG2a (Extended Data Fig. 8).

We observed that the 8E12-IgG2a mAbs had a much increased potency compared to the 8E12-IgG1 when used in passive transfer experiments (Fig. 4a), and titrating this antibody showed that three doses of 50 micrograms or more was sufficient to confer sterile protection (Fig.4b, Extended Data Fig. 9a). This demonstrated that recruitment of antibody-mediated immune effectors were important for parasite neutralisation and to quantify their relative contributions, we engineered three further mAbs which each lacked the binding sites for C1q (ΔC1q), FcRs (ΔFcR), or both (ΔC1qΔFcR)^22^ (Extended Data Fig. 8c). When used in passive protection experiments, we observed that mutation of the C1q binding site reversed the inhibition of parasite growth almost to the potency of the original IgG1 isotype, demonstrating that C1q-mediated complement recruitment was a major mechanism of immunological protection (Fig. 4c, Extended Data Fig. 9b). Mutating the FcR binding site also relieved the inhibition of parasite growth, but to a much lesser extent, while mutation of both C1q and FcR sites inhibited growth with a similar potency as the IgG1 isotype (Fig. 4c, Extended Data Fig. 9b). These experiments revealed that anti-IFX antibodies inhibited parasite multiplication by several immune mechanisms dominated by the recruitment of complement but also revealed roles for FcR binding, and direct binding and inhibition of IFX function.

**Figure 4.**
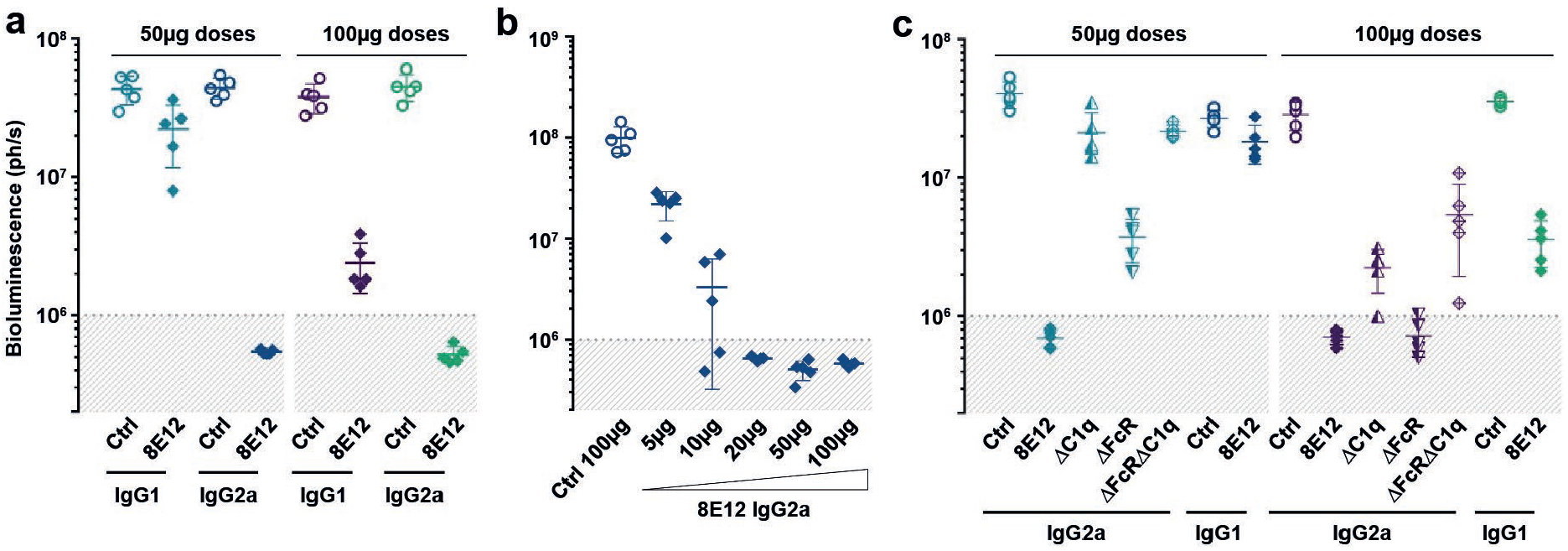
Multiple mechanisms of antibody-mediated anti-IFX immunological protection dominated by complement recruitment. **a,** Anti-IFX 8E12-IgG2a mAbs passively protect more potently against *T. vivax* infection than 8E12-IgG1 mAbs. **b**, Dose titration of 8E12-IgG2a mAb compared to isotype-matched control. **c,** Passive transfer of 8E12-IgG2a mAbs containing mutations that prevent binding to C1q (ΔC1q), FcRs (ΔFcR) or both (ΔFcRΔC1q) relieved the inhibition of parasite multiplication to differing degrees demonstrating multiple mechanisms of antibody-based immunological protection including a major role for complement. Parasitaemia was quantified at day 5 using bioluminescence, data points represent individual animals and grey shading indicates background bioluminescence; bars indicate mean ± SD. One of two independent experiments with very similar outcomes is shown.

### IFX is highly conserved across isolates

To further assess and develop IFX as a potential vaccine target, we tested appropriate routes of administration and other adjuvants that would bias antibody responses towards more protective isotypes. Of two selected adjuvants that have previously been used in veterinary vaccines and can be delivered subcutaneously, we found that the saponin-based adjuvant Quil-A elicited consistent antibody titres equivalent to the protective responses induced by alum, of which a large proportion were of the IgG2 isotypes (Extended Data Fig. 10); these mice were potently protected against parasite challenge (Fig. 5a). One potential challenge with subunit vaccines is that the genes encoding antigens eliciting protective immune responses in natural infections can be subject to diversifying selection, potentially leading to strain-specific immunity and thereby limiting the usefulness of the vaccine^23^. We therefore analysed the IFX gene sequence in 29 cosmopolitan *T. vivax* genomes and showed that it was highly conserved by comparison to other surface antigens. We observed only a single non-synonymous polymorphism in 2/29 isolates (Fig. 5b) and the frequency of the mutation in these two strains was very low (0.05), demonstrating that IFX is almost completely invariant across parasite strains. This high level of sequence conservation across isolates suggests that it is not a target of host immune responses and consistent with this, sera from naturally-infected cattle were not immunoreactive to IFX (Fig. 5c). Finally, a successful vaccine must be able to elicit long-lasting protection and so we repeatedly challenged IFX-vaccinated mice over 100 days after receiving their final immunisation. We observed that mice remained fully protected even with parasites that were delivered subcutaneously (Fig. 5d).

**Figure 5.**
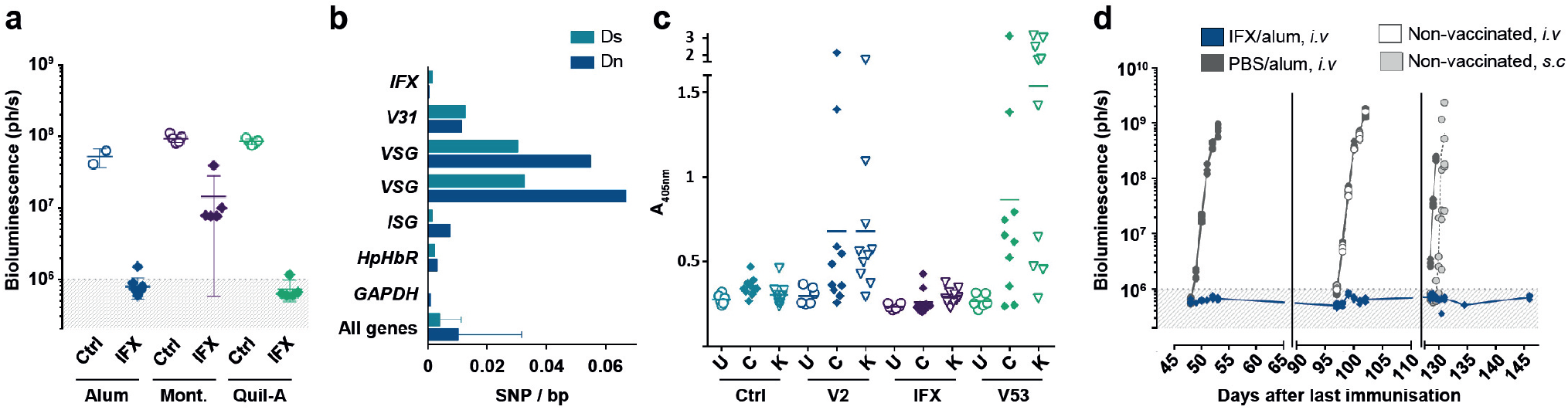
IFX is highly conserved and can elicit long-lasting immunity to *Trypanosoma vivax* infections. **a,** Comparing veterinary adjuvants using subcutaneous delivery demonstrates that Quil-A is as effective as IFX/alum positive control. Parasitaemia was quantified at day 5 using bioluminescence, data points represent individual animals and grey shading indicates background bioluminescence; bars indicate mean ± SD. **b,** Parasite population genetic analysis show *IFX* is highly conserved compared to other genes; mean synonymous (Ds) and nonsynonymous (Dn) substitution densities are shown + SD where appropriate. **c,** IFX is not immunogenic in the context of a natural infection. Immunoreactivity to the indicated proteins in sera from Cameroonian (C), Kenyan (K) or uninfected UK control cattle (U); bar indicates median. **d,** IFX/alum-vaccinated mice were protected from two intravenous and one final subcutaneous *T. vivax* challenge given over 100 days following the final booster immunisation.

## Discussion

We have shown that it is possible to elicit apparently sterile protective immunity to an experimental trypanosome infection by immunising with a recombinant subunit vaccine corresponding to the ectodomain of an invariant cell surface parasite protein termed IFX. The localisation of IFX to the flagellum membrane at the boundaries of where the flagellum is in contact with the parasite cell body suggests that it performs a role in flagellar structure and function. Our demonstration that antibodies are required for immunity raises important questions about the immunoprotective mechanisms employed by trypanosomes, and importantly, their vulnerabilities that can be exploited to develop vaccines. The inhibition of parasite growth by antibodies to IFX suggest that the *T. vivax* VSG surface coat cannot fully shield it from antibody binding, and that anti-IFX antibodies are not removed by endocytosis within the flagellar pocket from the parasite surface with sufficient rapidity to prevent antibody-mediated immune effector recruitment. The finding that the IFX gene sequence was highly conserved across parasite isolates and sera from infected cattle living in endemic regions were not immunoreactive to IFX suggest that natural parasite infections in some species can subvert host immunity to avoid eliciting protective antibody responses. These mechanisms could include perturbations of the B-cell compartment which have been described in experimental models of other trypanosome species^24–26^, or that the IFX protein may not be suitably presented to the host immune system in the context of a natural infection. The discovery of an antigen that can elicit protection to a trypanosome infection provides optimism and a technical roadmap that could be followed to identify vaccine antigens not just for other trypanosome species, but also parasites that have thus far proved intractable to vaccine development. Finally, IFX represents a very attractive vaccine candidate for an important livestock disease that has been a major barrier to the socioeconomic development of sub-Saharan Africa.

## Methods

### Mouse strains and ethical approvals

All animal experiments were performed under UK Home Office governmental regulations (project licence numbers PD3DA8D1F and P98FFE489) and European directive 2010/63/EU. Research was ethically approved by the Sanger Institute Animal Welfare and Ethical Review Board. The animals used in this study were female *Mus musculus* strain BALB/c which were obtained from a breeding colony at the Research Support Facility, Wellcome Sanger Institute.

### Cell lines and antibodies

Recombinant proteins and antibodies used in this study were expressed in HEK293E^27^ or HEK293-6E^28^ kindly provided by Yves Durocher (NRC, Montreal). Neither cell line was authenticated but were regularly tested for mycoplasma (Surrey Diagnostics, UK) and found to be negative. The antibodies used in this study are detailed below. Primary antibodies: six anti-IFX mouse monoclonal antibodies were selected and validated in this study: 6B3, 8C9, 3D12, 2H3, 10E2 and 8E12 from hybridomas all secreting IgG1 isotypes. A rabbit polyclonal antibody to the entire ectodomain of IFX was generated by Cambridge Research Biochemicals and validated by ELISA against the recombinant IFX ectodomain. The 8E12 antibody was cloned and expressed recombinantly as a mouse IgG2a isotype as described below. Mouse isotype control antibodies were: IgG1 (MOPC-21, BioXcell) and IgG2a (C1.18.4, BioXcell). Antibodies used for *in vivo* leukocyte cell depletion were: anti-mouse Cd4 (clone GK1.5, BioXcell), anti-mouse Cd8 (clone 2.43, BioXcell), anti-mouse Nk1.1 (clone PK136, BioXcell), and control anti-keyhole limpet hemocyanin (clone LTF-2, BioXcell). Antibodies used for protein quantification for ELISAs were mouse monoclonal anti-His (His-Tag mAb, EMD-Millipore), and biotinylated mouse anti-rat Cd4 (clone OX68). Secondary antibodies used were goat anti-mouse alkaline phosphatase conjugated secondary (Bethyl) and rabbit anti-bovine alkaline phosphatase conjugated secondary (Sigma). Mouse antibody isotypes were determined using the mouse monoclonal antibody isotyping kit (Sigma).

### Vaccine target identification and expression

The *T. vivax* genome was searched for proteins encoding predicted type I, GPI-anchored and secreted proteins using protein feature searching in TriTrypDB^29^. The regions corresponding to the entire predicted extracellular domains of *T. vivax* cell-surface and secreted proteins from the Y486 strain were determined by using transmembrane^30^ and GPI-anchor^31^ or signal peptide^32^ prediction software. Protein sequences encoding the predicted extracellular domain and lacking their signal peptide, were codon-optimized for expression in human cells and made by gene synthesis (GeneartAG, Germany and Twist Bioscience, USA). The sequences were flanked by unique NotI and AscI restriction enzyme sites and cloned into a pTT3-based mammalian expression vector^27^ between an N-terminal signal peptide to direct protein secretion and a C-terminal tag that included a protein sequence that could be enzymatically biotinylated by the BirA protein-biotin ligase^33^ and a 6-his tag for purification^34^. The ectodomains were expressed as soluble recombinant proteins in HEK293 cells as described^35,36^. To prepare purified proteins for immunisation, between 50 mL and 1.2 L (depending on the level at which the protein was expressed) of spent culture media containing the secreted ectodomain was harvested from transfected cells, filtered and purified by Ni^2+^ immobilised metal ion affinity chromatography using HisTRAP columns using an AKTAPure instrument (GEHealthcare, UK). Proteins were eluted in 400 mM imidazole as described^37^, and extensively dialysed into HBS before being quantified by spectrophotometry at 280 nm. Protein purity was determined by resolving one to two micrograms of purified protein by SDS-PAGE using NuPAGE 4–12 % Bis Tris precast gels (ThermoFisher) for 50 minutes at 200 V. Where reducing conditions were required, NuPAGE reducing agent and anti-oxidant (Invitrogen) were added to the sample and the running buffer, respectively. The gels were stained with InstantBlue (Expedeon) and imaged using a c600 Ultimate Western System (Azure biosystems). Purified proteins were aliquoted and stored frozen at −20 °C until use. Where enzymatically monobiotinylated proteins were required to determine antibody titres by ELISA, proteins were co-transfected with a secreted version of the protein biotin ligase (BirA) as described^36^, and extensively dialysed against HEPES-buffered saline and their level of expression determined by ELISA using a mouse monoclonal anti-His antibody (His-Tag mAb, EMD Millipore) as primary antibody and a goat anti-mouse alkaline phosphatase-conjugated secondary (Bethyl).

### Vaccine formulation and administration

For the initial screening of antigens, aliquots of purified protein for immunisation were thawed, diluted, and mixed 50 % v/v with alhydrogel adjuvant 2 % (InvivoGen) for two hours at room temperature. For each antigen, groups of five 6 to 8-week old female BALB/c mice were immunised intraperitoneally using a prime and two boosts strategy using the amounts of protein documented in Extended Data Table 1. For retesting those antigens which had shown some effect in the preliminary screen, one group of 15 animals received three intraperitoneal immunisations of the query protein adjuvanted in alum using similar amounts as used in the initial screen (Extended Data Table 1); a control group, also 15 animals, received the adjuvant alone. For evaluating different IFX vaccine adjuvant formulations, groups of five mice received three immunisations of 50 μg IFX adjuvanted with either alhydrogel, Montanide ISA 201 VG, or Quil-A in a total volume of 200 μL. IFX was formulated with Montanide ISA 201 VG according to the manufacturer’s instructions using a stirrer to create the water-in-oil emulsion. IFX was mixed in a 1:1 (v/v) ratio with Quil-A adjuvant using a 0.5 mg mL^−1^ solution. IFX/Montanide and IFX/Quil-A vaccines were administered subcutaneously at two different injection sites (100 μL per site), and IFX/alhydrogel formulation was administered intraperitoneally.

### Quantification of serum antibody titres by ELISA

To determine the serum antibody responses to immunised proteins, blood biopsies were collected between ten to twelve days after the final immunisation from the tail of each animal and clotted for two hours at room temperature. Cells were removed by centrifugation, the serum collected, supplemented with sodium azide to a final concentration of 2 mM as a preservative and stored at −20 °C until use. Cattle sera were donated from archived material at the University of Liverpool, originally collected from natural *T. vivax* infections in Cameroon (Northwest state), and Kenya (Western state) where the infection was positively identified by thick blood smear and the parasite identified as *T. vivax* using the VerY Diag field test. To determine the antibody titre against an antigen of interest, individual sera were initially diluted 1:1000 and then six four-fold serial dilutions in PBST/2 % BSA were prepared. These dilutions were pre-incubated overnight at room temperature with 100 μg mL^−1^ of purified rat Cd4d3+4-BLH protein to adsorb any anti-biotin/his tag antibodies. Sera were transferred to streptavidin-coated ELISA plates on which the biotinylated target antigen was immobilised. To ensure that all anti-tag antibodies were adsorbed, binding of the lowest dilution of antisera was also tested against biotinylated rat Cd4d3+4-BLH protein similarly immobilised on the ELISA plate to confirm the absence of any anti-tag immunoreactivity^38^. Sera were incubated for one hour at room temperature followed by three washes with PBST before incubating with an anti-mouse IgG secondary antibody conjugated to alkaline phosphatase (Sigma) used as a 1:5000 dilution for one hour. Following three further washes with PBST, 100 μL of 1 mg mL^−1^ Sigma 104 phosphatase substrate was added and substrate hydrolysis quantified at 405 nm using a plate reader (Spark, Tecan). To quantify immunoreactivity to *T. vivax* antigens in the context of natural infections, cattle sera were diluted 1:800 in PBST/2 % BSA and incubated for two hours at room temperature with biotinylated ectodomains of V2, V53, IFX or control rat Cd200, adsorbed on the microtiter plate. Following three washes with PBST, a secondary rabbit anti-bovine IgG antibody (Sigma) diluted 1:20,000 was incubated for one hour and washed three times with PBST before adding colourimetric phosphatase substrate and acquiring absorbance readings as described above.

### Antibody isotyping

Isotyping of the monoclonal antibodies and polyclonal sera responses was performed using the Mouse Monoclonal Antibody Isotyping Kit (Sigma-Aldrich), according to the manufacturer’s instructions. Briefly, the biotinylated ectodomain of the IFX protein was immobilised on a streptavidin-coated plate, incubated with sera diluted 1:1000 in PBST/2 % BSA or hybridoma supernatants, washed in PBST before adding isotype-specific goat anti-mouse secondary antibodies diluted 1:1000. Binding was quantified with an alkaline-phosphatase-conjugated rabbit anti-goat tertiary antibody (1:5000, Sigma) followed by a colourimetric phosphatase substrate, and hydrolysis products quantified by absorbance readings at 405 nm.

### *Trypanosoma* parasite strain and maintenance

A transgenic form of *Trypanosoma vivax* genetically engineered to ubiquitously express the firefly luciferase enzyme^14^ was kindly provided by Paola Minoprio, Institut Pasteur, Paris. The parental strain of this parasite is the IL1392 line derived from the Y486 strain used for genome sequencing^16^ and is fully documented by Chamond *et al*.^15^. Parasites were initially recovered from a frozen stabilate by intraperitoneal administration into two BALB/c female mice. Parasites were maintained by weekly serial blood passage in wild type female BALB/c mice by taking a blood biopsy, quantifying living parasites in PBS/20 mM D-glucose by microscopy and infecting four naive mice intravenously. During the course of the project, two further aliquots of frozen parasites were thawed and then used for infection challenges, no significant differences in the kinetics of infection were observed. Luciferase-expressing *T. congolense* parasites were a kind gift from Bill Wickstead and Caterina Gadelha, University of Nottingham, and maintained by weekly serial intravenous blood passage in wild type female BALB/c mice.

### *Trypanosoma vivax* infections

For infection challenges, bloodstream forms of *T. vivax* parasites were obtained from the blood of an infected donor mouse at the peak of parasitaemia, diluted in PBS/20 mM D-glucose, quantified by microscopy and used to infect mice by intravenous injection. While establishing the infection model in our facility, we observed that the *T. vivax* parasite was labile and gradually lost virulence once removed from living mice. To reduce the possibility of any artefactual protective effects being due to the loss of parasite virulence during the challenge procedure, we screened the protective effects of antigens in a cohort design. Each cohort contained six cages of five animals: four cages contained mice immunised with a different query subunit vaccine candidate, and the other two cages contained control mice immunised with adjuvant alone. Vaccinated animals were rested for four to eight weeks after the final immunisation to mitigate any possible non-specific protective effects elicited by the adjuvant. During the infection procedure, the mice in the control cages were challenged first and last, and the data from the cohort only used if the infections in the control mice from the two cages were comparable. During the infection procedures, parasites were outside of a living mouse for no more than 40 minutes. Mice were normally challenged by intravenous delivery of 10^2^ (cohorts 1-7, 10-11) to 10^3^ (cohorts 8 and 9) parasites for the initial screening and passive transfer protection experiments, but were also challenged intraperitoneally during the establishment of the model and subcutaneously when investigating the duration of protection. The animals were not randomised between cages and the operator was not blinded to the group condition. Occasionally, individual infected mice within a group unexpectedly exhibited only background levels of bioluminescence which was attributed to the injected luciferin substrate not distributing from the site of delivery, possibly due to mislocalization of the injection bolus; in these instances, these animals were excluded from the analysis. Groups were compared using bioluminescence quantification as a proxy for parasitaemia and one-way ANOVA with Dunnett’s post-hoc test unless specified.

### Quantification of *Trypanosoma vivax* infections by bioluminescent *in vivo* imaging

The luciferase substrate D-luciferin (potassium salt, Source BioScience, UK) was reconstituted to 30 mg mL^−1^ in Dulbecco’s PBS (Hyclone), filter-sterilised (0.22 μm) and stored in aliquots at −20 °C. Aliquots were thawed and administered to animals at a dose of 200 mg kg^−1^, by intraperitoneal injection ten minutes before bioluminescence acquisitions. The mice were given three minutes of free movement before being anaesthetized with 3% isoflurane and placed in the imaging chamber where anaesthesia was maintained for acquisition. An average background bioluminescence measurement was determined by luciferin administration in five female BALB/c mice and calculating the mean whole-body bioluminescence; where appropriate, this value is indicated as a light grey shading on bioluminescence plots. To determine long-term persistence of the parasites in different organs of infected mice, animals were administered with luciferin, imaged, and then euthanised with an overdose of anaesthetic. Mice were then perfused with PBS until the perfusion fluid ran clear, the organs dissected, arranged on a petri dish, and bathed in PBS containing 20 mM glucose and 3.3 mg mL^−1^ luciferin for imaging. Emitted photons were acquired by a charge coupled device (CCD) camera (IVIS Spectrum Imaging System, Perkin Elmer). Regions of interest (ROIs) were drawn and total photons emitted from the image of each mouse were quantified using Living Image software (Xenogen Corporation, Almeda, California), the results were expressed as the number of photons sec^−1^. Where necessary, peripheral parasitaemia was quantified by direct microscopic observation as previously described^15^. Briefly, five microliters of blood obtained from the tail vein were appropriately diluted in PBS containing 20 mM glucose and parasite counts were expressed as number of parasites per blood milliliter.

### Passive transfer of immunity

To obtain sufficient sera for adoptive transfer experiments, fifty 6 to 8-week-old female BALB/c mice were immunised intraperitoneally three times with 20 μg of purified IFX adjuvanted in alum, with each immunisation separated by two weeks. Nine days after the final immunisation, sera were collected as above, aliquoted, and stored at −20 °C until use. For passive transfer experiments, groups of six to eight-week-old female BALB/c mice were dosed three times with either sera or purified monoclonal antibodies on three consecutive days; three hours after the second dosing, mice were challenged intravenously with 10^2^ *T. vivax* parasites. When using immune serum for passive transfer protection experiments, doses of 100 and 200 μL of sera from either IFX-vaccinated mice or non-immunised control mice were administered. For monoclonal antibodies, the purified antibody was diluted to the required dose in PBS and 200 μL administered intravenously. Control isotypes antibodies used were MOPC-21 for the IgG1 isotype and C1.18.4 for the IgG2a isotype (both from BioXcell).The serum half-life for mouse IgG1 and IgG2a are known to be between 6 to 8 days^39^.

### *In vivo* cell depletion

Groups of five mice were immunised three times with 50 μg doses of purified IFX to induce protective immunity to *T. vivax*. To deplete immune animals of defined leucocyte lineages, animals within each group were depleted by intra-peritoneal administration of lineage-specific monoclonal antibodies using standard procedures. Briefly, NK cells were depleted by four injections of 500 μg of the PK136 mAb that targets the Nk1.1 glycoprotein at days −5, −1, 0 and 2 post-infection. Mouse CD4 and CD8 T-lymphocytes were depleted by one intraperitoneal 750 μg injection of the mAbs targeting Cd4 (clone GK1.5) or Cd8 (clone 2.43) receptors, respectively, the day prior of the infection. The LFT-2 mAb (750 μg) was used as an isotype-matched control antibody. Mice were challenged with 10^2^ *T. vivax* parasites and parasitaemia quantified using bioluminescent imaging as described.

### Trypanosome genomic sequence analysis

To identify if IFX had any homologues in other *Trypanosome* species, the entire IFX sequence was analysed with Interproscan which showed that it does not contain any known protein domains, other than the predicted N-terminal signal peptide and transmembrane helix. Comparison of the predicted IFX protein sequence with all the other sequenced *Trypanosoma* spp. genomes in TriTrypDB^29^ using tBLASTx returned no significant matches; moreover, comparison of a Hidden Markov Model of the IFX protein sequence with all *T. brucei, T. congolense* and *T. cruzi* proteins using HMMER also produced no matches demonstrating IFX is unique to *T. vivax*. To confirm that IFX is present in a single-copy, the IFX protein sequence was compared with a six-frame translation of the genome sequence using tBLASTn to identify any sequence copies (annotated or not) with >98% amino acid identity typical of allelic variation.

Illumina sequencing reads from 29 clinical strains isolated from Nigeria, Togo, Burkina Faso, The Gambia, Ivory Coast, Uganda and Brazil were mapped to the *T. vivax* Y486 reference sequence using BWA^40^ before SNPs were called using the GATK4 analysis toolkit^41^. Indels and variant positions with QD < 2.0, FS > 60.0, MQ < 40.0, MQRankSum < −12.5 or ReadPosRankSum < −8.0 were excluded to produce a final list of 403,190 SNPs. Individuals were classified as missing if allele calls were supported by fewer than 3 reads. Coding SNPs and synonymous/non-synonymous codon alterations were identified by comparison to the reference annotation using a custom Biopython script; pi-values were calculated on a per site basis using vcftools^42^. Genes selected for comparison were *V31* (TvY486_0003730), two *VSG*s (TvY486_0031620, TvY486_0040490), *ISG* (TvY486_0503980), *HpHbR* (TvY486_0040690) and the “housekeeping” gene *GAPDH* (TvY486_1006840).

### Electron microscopy

*T. vivax* parasites were resuspended in 1 % paraformaldehyde in PBS for 30 minutes (all steps at room temperature), washed three times in PBS, blocked with PBS/glycine followed by 5 % foetal calf serum for 30 minutes and then incubated with a mouse monoclonal antibody to IFX (clone 8E12) for 1 hour. After rinsing, the parasites were incubated with goat anti-mouse IgG preadsorbed to 10 nm gold particles (ab27241 Abcam) for 30 minutes, washed, and fixed in a mixture of 2 % paraformaldehyde and 2.5 % glutaraldehyde in 0.1 M sodium cacodylate buffer for 30 minutes. After washing again, the parasites were post-fixed in 1 % osmium tetroxide for 30 minutes, dehydrated in an ethanol series, embedded in epoxy resin and 60 nm ultrathin sections were cut on a Leica UC6 ultramicrotome, contrasted with uranyl acetate and lead citrate and examined on a 120 kV FEI Spirit Biotwin using a Tietz F4.16 CCD camera. The density of anti-IFX gold particle staining was determined by counting the number of gold particles per micron of membrane on both sagittal and transverse sections. Membrane lengths were determined using the segmented line function in ImageJ and known image scaling factor. To assign dorsal and ventral sectors, a line was drawn across (transverse) or along (sagittal) the flagellum midpoint.

### Anti-IFX antibody selection and characterisation

To raise polyclonal antisera against IFX, the entire ectodomain of IFX was expressed and purified and injected into rabbits (Cambridge Research Biochemicals, Billingham, UK). The sera were purified on Hi-Trap Protein G HP columns (GE Healthcare) according to the manufacturer’s instructions. Hybridomas secreting monoclonal antibodies to IFX were selected using standard protocols as described^43^. In brief, the SP2/0 myeloma cell line was grown in advanced DMEM/F12 medium (Invitrogen, CA, USA) supplemented with 20 % fetal bovine serum, penicillin (100 U mL^−1^), streptomycin (100 μg mL^−1^) and L-glutamine (2 mM). Following spleen dissection and dissociation, 10^8^ splenocytes were fused to 10^7^ SP2/0 myeloma in 50 % PEG (PEG 1500, Roche, Hertfordshire, UK), using standard procedures. The resulting hybridomas were plated over ten 96-well plates and initially grown in advanced DMEM/F12 medium (Invitrogen) supplemented with 20 % fetal bovine serum, penicillin (100 U mL^−1^), streptomycin (100 μg mL^−1^) and L-glutamine (2 mM) before addition of hypoxanthine-aminopterin-thymidine (HAT) selection medium 24 hours after the fusion. After 11 days, hybridoma supernatants were harvested to determine the presence of antibodies reacting to the IFX protein using an ELISA-based method as previously described^43^. Six wells (2H3, 3D12, 6B3, 8C9, 8E12, and 8F10) containing hybridoma colonies secreting antibodies that reacted with IFX but not a control protein containing the same purification tags were identified and cultured for a further four days in HAT-selection medium. Hybridoma cells from each of the positive wells were cloned by limiting dilution over two 96-well plates at a density of 0.5 cells per well and grown in HAT-free SP2/0 conditioned medium. Eleven days later, twelve wells corresponding to each of the seven clones were selected and tested again by ELISA for reactivity to the IFX protein; three positive wells per clone were chosen for a second round of dilution cloning in the conditions described above. After a final test for reactivity to IFX, a single well from each of the seven positive clones was expanded and adapted to grow in Hybridoma-SFM serum-free medium (Thermo Fisher).

To determine the location of the anti-IFX monoclonal antibody epitopes, subfragments of the IFX ectodomain corresponding to the boundaries of predicted secondary structure (M1-T251, M1-S472, S135-T251 and N442-S535) were designed, produced by gene synthesis and cloned into a mammalian expression plasmid with an enzymatically biotinylated C-terminal tag (Twist Biosciences, USA). Biotinylated proteins were expressed as secreted recombinant proteins in HEK293 cells as described above and dialysed to remove free D-biotin. Biotinylated IFX fragments were immobilised on a streptavidin-coated plate and binding of the six mouse monoclonal antibodies was tested by ELISA and detected with an alkaline-phosphatase-conjugated anti-mouse secondary antibody (Sigma-Aldrich) as previously described^43^. Binding of a rabbit polyclonal antibody raised to the entire ectodomains of IFX (Cambridge Research Biochemicals) was used as a positive control for each of the subdomains, and detected with an alkaline-phosphatase-conjugated anti-rabbit secondary antibody (Jackson Immunoresearch).

For affinity-purification of monoclonal antibodies from hybridoma culture supernatants, spent supernatants were supplemented with 0.1 M sodium acetate, pH 5.0 immediately before purification on a HiTrap Protein G HP 1 mL column (GE Healthcare) using an AKTA pure instrument. Elution was performed in 0.1 M glycine, pH 2.7 followed by immediate neutralisation with 1 M Tris-HCl, pH 9.0. Purified antibodies were extensively dialysed against PBS and stored at 4 °C until use. To capture antibodies on streptavidin-coated sensor chips for biophysical interaction analysis, 300 μg of purified monoclonal antibodies were chemically biotinylated using a 20-fold molar excess of sulfo-NHS-biotin (ThermoFisher) for two hours at room temperature; to remove excess biotin the solutions were dialysed against 5 L PBS for 16 hours.

### Antibody affinity by surface plasmon resonance

Antibody affinities were determined by SPR essentially as described^44^ using a Biacore 8K instrument (GE Healthcare, Chicago, IL). To measure antibody interaction affinity rather than avidity, between 400 to 600 RU of biotinylated anti-IFX monoclonal antibodies were immobilised on a streptavidin-coated sensor chip prepared using the Biotin CAPture kit (GE Healthcare); a biotinylated mouse monoclonal antibody (OX68) was used as a non-binding control in the reference flow cell. The entire ectodomain of IFX was used as the analyte which was first purified and resolved by size exclusion chromatography on a Superdex 200 Increase 10/300 column (GE Healthcare, Chicago, IL) in HBS-EP (10 mM HEPES, 150 mM NaCl, 3 mM EDTA, 0.05 % v/v P20 surfactant) just prior to use in SPR experiments to remove any protein aggregates that might influence kinetic measurements. Increasing concentrations of two-fold dilutions of the entire ectodomain of IFX as a soluble analyte were injected at 30 μL min^−1^ for a contact time of 120 s and dissociation of 600 s. Both kinetic and equilibrium binding data were analyzed in the manufacturer’s Biacore 8K evaluation software version 1.1 (GE Healthcare, Chicago, IL). All experiments were performed at 37 °C in HBS-EP.

### Antibody cloning, isotype switching, mutagenesis, and purification

To switch the isotype of the 8E12 anti-IFX monoclonal antibody from IgG1, it was first necessary to amplify the genes encoding the rearranged light and heavy variable regions from the hybridoma; this was performed essentially as described^43^. Briefly, total RNAs were extracted from the cloned 8E12 hybridoma using the RNAqueous-micro total RNA isolation kit (Ambion) followed by reverse transcription with Superscript III (Thermo Fisher). PCR products encoding the rearranged heavy and light chain regions were individually amplified using sets of degenerate oligonucleotides and then assembled in a subsequent fusion PCR using a linker fragment to create a single PCR product containing both the rearranged light and heavy chains, as previously described^45^. The fusion PCR product was ligated using the NotI and AscI restriction sites into an expression plasmid obtained from Addgene (plasmid # 114561) in frame with the mouse constant IgG2a heavy chain^46^. Competent *E.coli* were transformed and purified plasmids used in small-scale transfections of HEK293 cells to identify those plasmids encoding functional antibodies as described^47^.

To perturb the recruitment of immune effectors in the murine IgG2a recombinant antibody and retain serum half-life, we mutated the C1q and FcR binding sites in the IgG2a constant heavy chain by site-directed mutagenesis as described^22^. Mutation to the binding site of Fcγ receptors (ΔFcR) was achieved by introducing the L234A and L235A substitutions using primers FcR_f_ −5’ GCACCTAACGCTGCAGGTGGACCATCCG 3’ and FcR_r_ - 5’ TGGTCCACCTGCAGCGTTAGGTGCTGGGC 3’. To abrogate C1q binding (ΔC1q), a single amino-acid change P329A was introduced using primers C1q_f_ – 5’ CAAAGACCTCGCTGCGCCCATCGAGAGAACC 3’ and C1q_r_ – 5’ GATGGCGCAGCGAGGTCTTTGTTGTTGACC 3’. In both cases, antibody mutagenesis was achieved by first amplifying 20 ng of an expression vector containing the mouse constant IgG2a heavy chain with each oligonucleotide separately for nine cycles (denaturation for 45 seconds at 94 °C; annealing for 40 seconds at 58 °C; elongation for 7 minutes and 30 seconds at 72 °C), using the KOD Hot Start DNA polymerase (Merck). Amplification reactions performed with complementary oligonucleotides were then mixed, 0.5 μL KOD Hot Start DNA polymerase was added to the reaction, and the amplification was resumed for a further 18 cycles. At the end of the reaction, half of the PCR reaction was digested with 20 U DpnI enzyme (New England Biolabs), which specifically cleaves methylated strands from the parental plasmid, for 3 hours at 37 °C before transforming 5 μL into TOP 10 chemically-competent bacteria (Invitrogen). Mutations were confirmed in selected clones by DNA sequencing. To generate a double mutant lacking both the C1q and FcR binding sites (ΔC1qΔFcR), site-directed mutagenesis was performed as described above on an expression plasmid containing the FcR mutation, using the set of oligonucleotides designed for C1q mutagenesis. Both single mutants and the double mutant backbones were doubly digested with NotI and AscI restriction enzymes and the fusion PCR product encoding the variable regions of the 8E12 recombinant antibody described above cloned into them, plasmids purified and verified by sequencing.

Antibodies were produced by transfecting HEK293 cells with plasmids encoding the recombinant 8E12-IgG2a monoclonal antibody with the wild-type IgG2a heavy chain, single mutants that lacked C1q and FcR binding, and the double mutant. Six days after transfection, the cell culture supernatant was harvested and the recombinant antibodies were purified on a HiTrap Protein G HP 1mL column, according to the manufacturer’s instructions as described^43^.

### Data availability

All data generated or analysed during this study are included in this published article and/or available from the corresponding author on reasonable request.

## Acknowledgements

This research was funded by the Wellcome Trust (grant 206194) and BBSRC (grant BB/S001980/1). A. R. R. was supported by a PhD studentship funded by FONDECYT-CONCYTEC, the National Council of Science, Technology and Innovation from Peru (grant contract number 001-2016-FONDECYT). We are indebted to Institut Pasteur for providing us with the *T. vivax* luciferase line. We are grateful to Sanger Institute animal technicians for their support with animal work, Natasha Karp for advice on experimental design, Craig Mackenzie for preliminary work on anti-IFX mAbs, Eve Coomber for help with hybridoma tissue culture, and Mark Carrington and Tim Rowan for helpful discussions.

## Author contributions

D. A. and G.J.W. designed the study. D.A. prepared proteins, and performed immunisations and parasite challenges with help from S.C., C.B., H.O. and K.H. C.C. selected hybridomas, cloned the 8E12 mAb, did the isotype switching, mutagenesis and purification. D.A.G. performed immunogold labelling. C. T. generated hybridomas. F.G. and M.K. performed SPR experiments and analysis. A. R. R., C. W. D. and A. P. J. provided and analysed *T. vivax* genome sequences and provided infected cattle sera. D.A. and G.J.W. prepared the manuscript with comments from all co-authors.

