## Supplemental Information for "An invariant *Trypanosoma vivax* vaccine antigen inducing protective immunity"

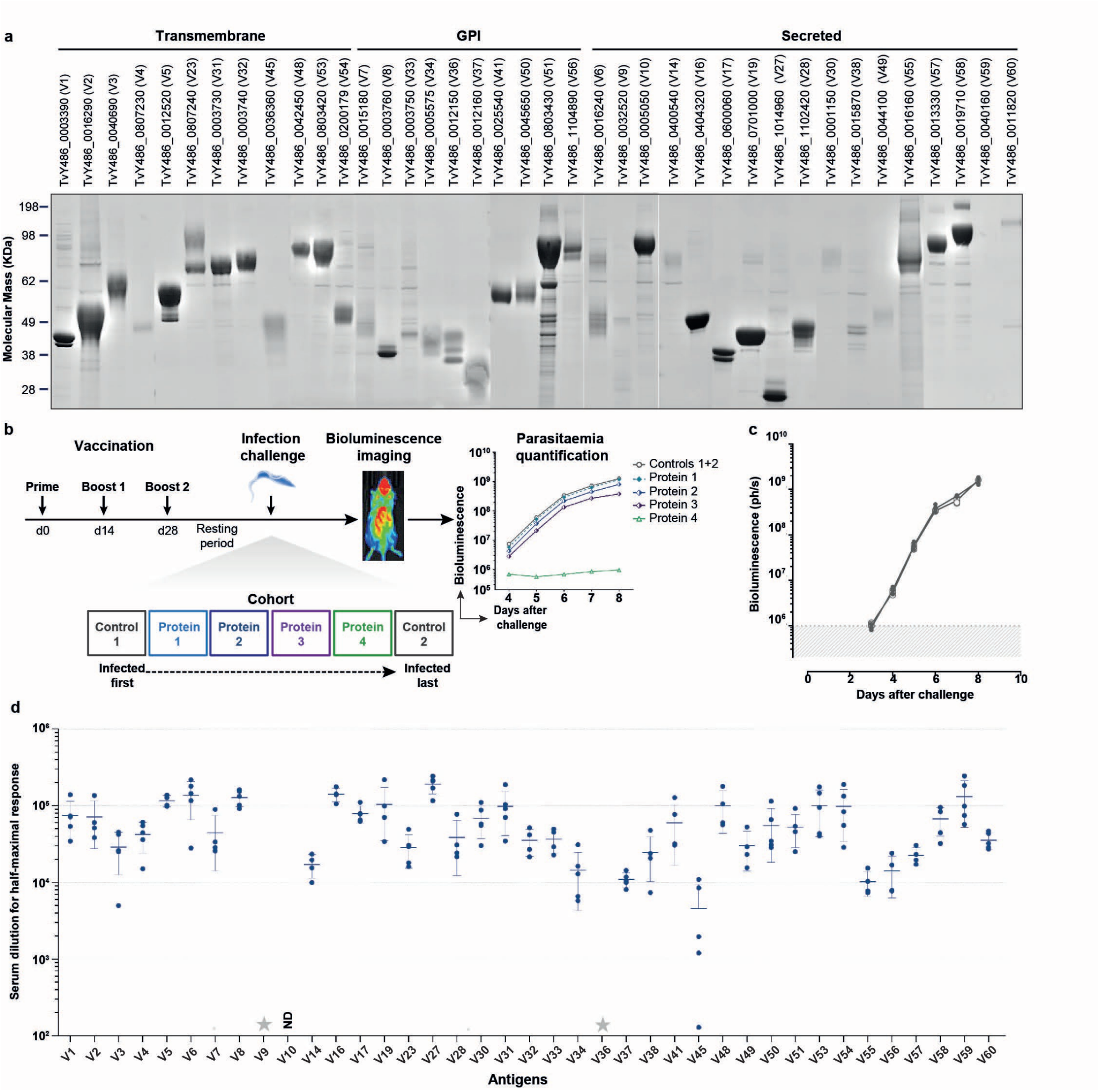


**Extended Data Figure 1 | *T. vivax* vaccine candidate antigens, organisation of the protection screen and antibody titres. a,** Vaccine candidates were expressed as soluble recombinant proteins in HEK293 cells, purified, and resolved by SDS-PAGE to determine protein integrity and purity. **b,** Mice were vaccinated with a protein-in-alum formulation using a prime and two boost regime and rested before challenge with the luciferase-expressing *T. vivax* parasite line and parasitaemia quantified using bioluminescent imaging. Vaccine candidates were tested in cohorts containing two control cages which were infected first and last to ensure any effect on the reduction of parasite multiplication was not confounded by the loss of parasite virulence. **c**, Parasite multiplication was identical in animals treated with adjuvant alone compared to naive mice. A group of five mice were immunised three times with alum alone (filled circles) and rested for 6 weeks and the infection with bioluminescent *T. vivax* was compared to naive mice (open circles); grey shading indicates background bioluminescence. **d,** Serum dilutions for the half-maximal responses for each antigen. Data points are individual mice and bars represent mean ± SD; $n = 5$. ND = not determined, and grey stars indicate no detectable response.

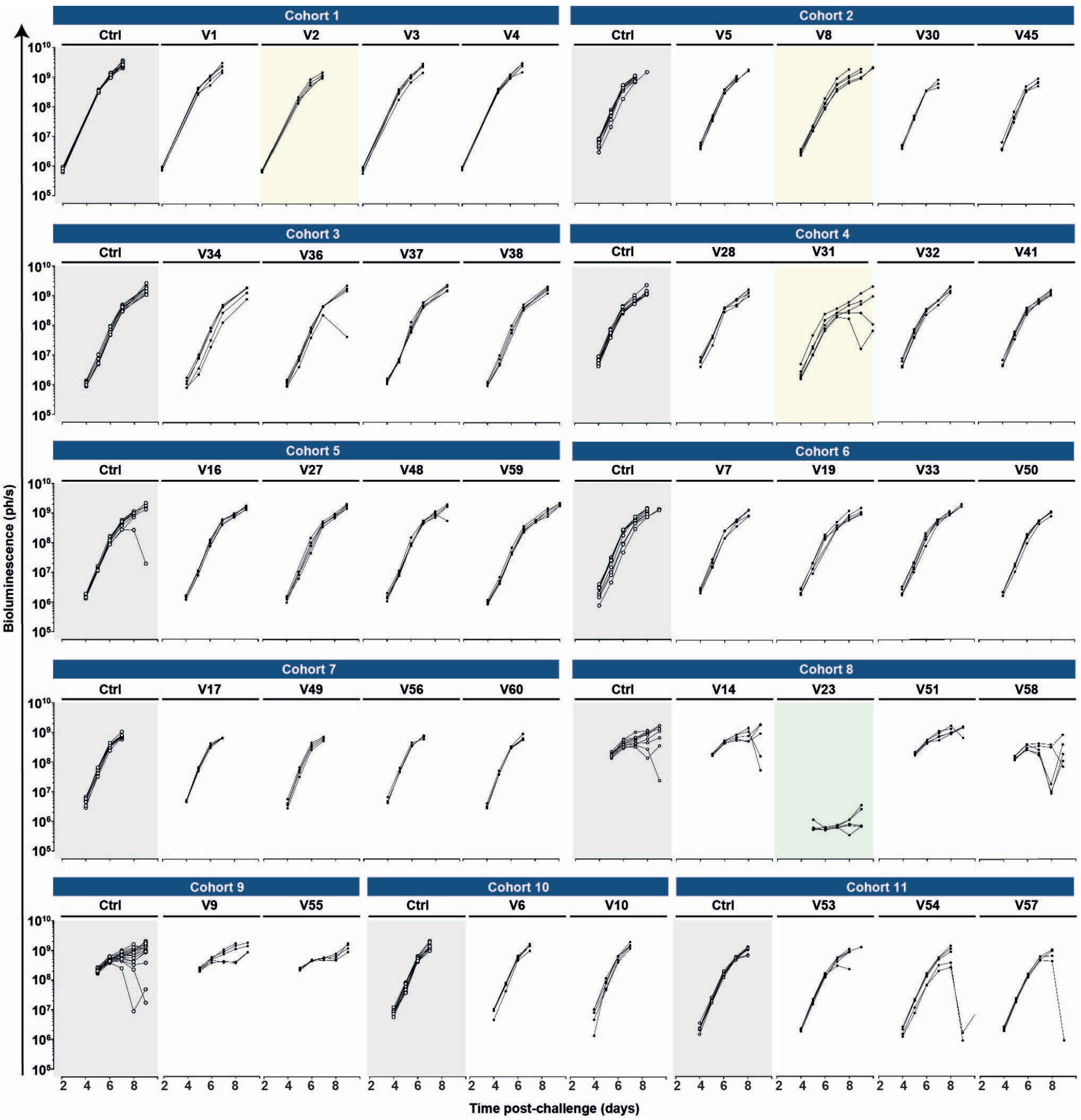


**Extended Data Figure 2 | Summary of systematic genome-led reverse vaccinology screen that identified subunit vaccine candidates for *Trypanosoma vivax*.** Bioluminescence is used as a proxy for parasitaemia and is shown for each mouse in the indicated days post-challenge. Candidates are identified using their "V number" and organised into their screening cohorts. The two cages of adjuvant-only controls for each cohort are highlighted in grey. The majority of candidates had no effect on the ascending phase of parasitaemia, and are left unshaded. Three candidates (V2, V8, V31) that had a statistically significant effect on infection are highlighted in pale yellow, and the candidate that elicited strong protection (V23) is highlighted in pale green. Data points are bioluminescence readings from individual animals. The occasional reductions in parasitaemia followed by rebound after day 8 are likely to be due to protective anti-VSG responses and selection of an antigenically distinct variant.

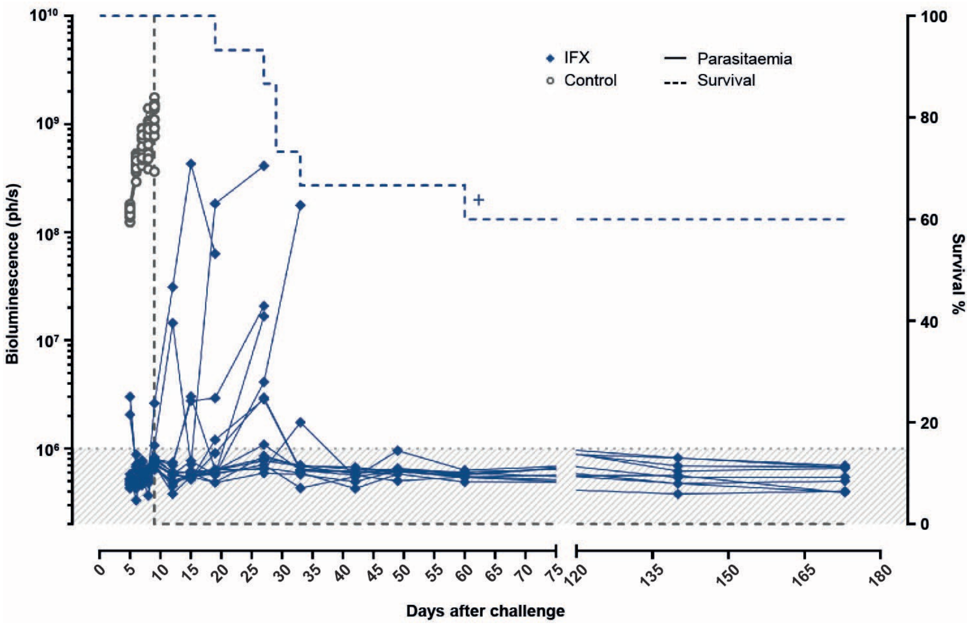


**Extended Data Figure 3 | Replication of strong protective effects in a larger group of IFX/V23-vaccinated mice.** Fifteen mice were immunized with purified soluble IFX/V23 recombinant protein adjuvanted in alum and challenged with transgenic luciferase-expressing *T. vivax*. Parasitaemia was quantified on the indicated days after parasite challenge using bioluminescence; controls are a cohort of 15 animals treated with adjuvant only. Ten out of the 15 mice were protected until at least day 170. Data points represent individual animals and grey shading indicates bioluminescence thresholds of uninfected mice. Where animals had to be euthanized for health reasons thought to be unrelated to the infection, this is indicated by a cross.

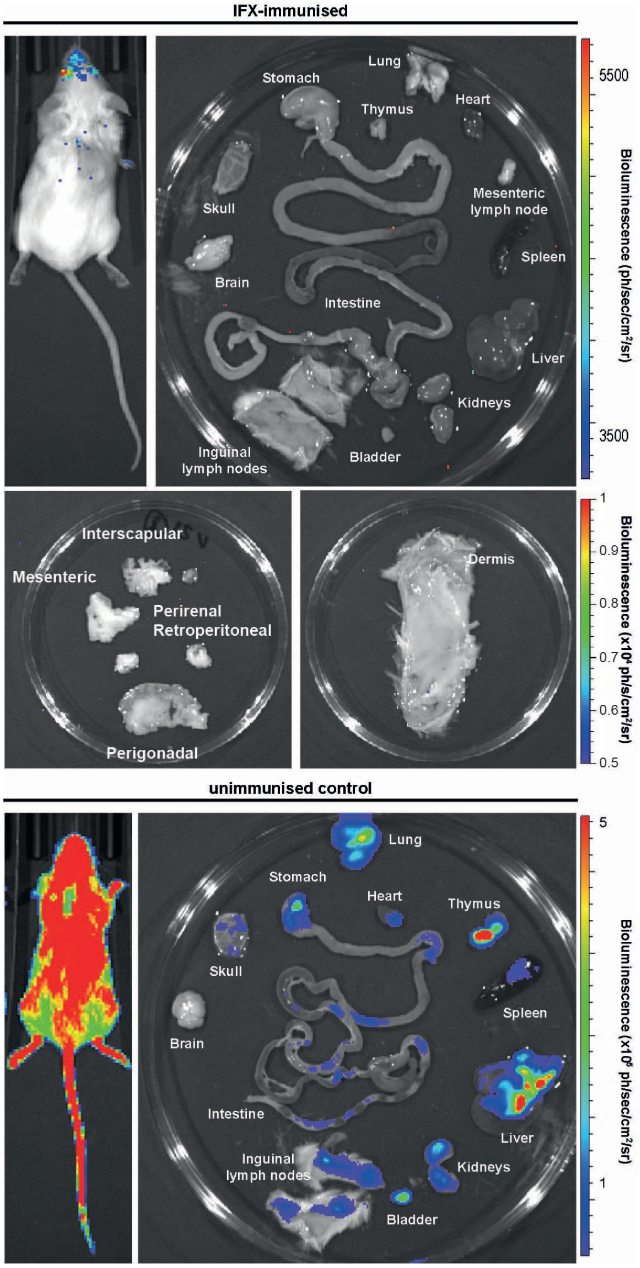


**Extended Data Figure 4 | Parasites do not detectably persist in the organs, adipose tissue, or dermis of IFX/V23-vaccinated *Trypanosoma vivax*-challenged mice.** An IFX/V23-immunised mouse was challenged with luciferase-expressing transgenic *T. vivax* parasites and protection from infection relative to controls was established. The animal was rested and nine months later, injected with luciferin to detect residual parasites using bioluminescence. No bioluminescent signals above background were detected in vaccinated animals either when the whole animal was imaged, or within the dissected organs, adipose deposits and dermis of IFX/V23-immunised (top panels). An unimmunised animal was used 8 days after parasite challenge as a positive control (lower panels).

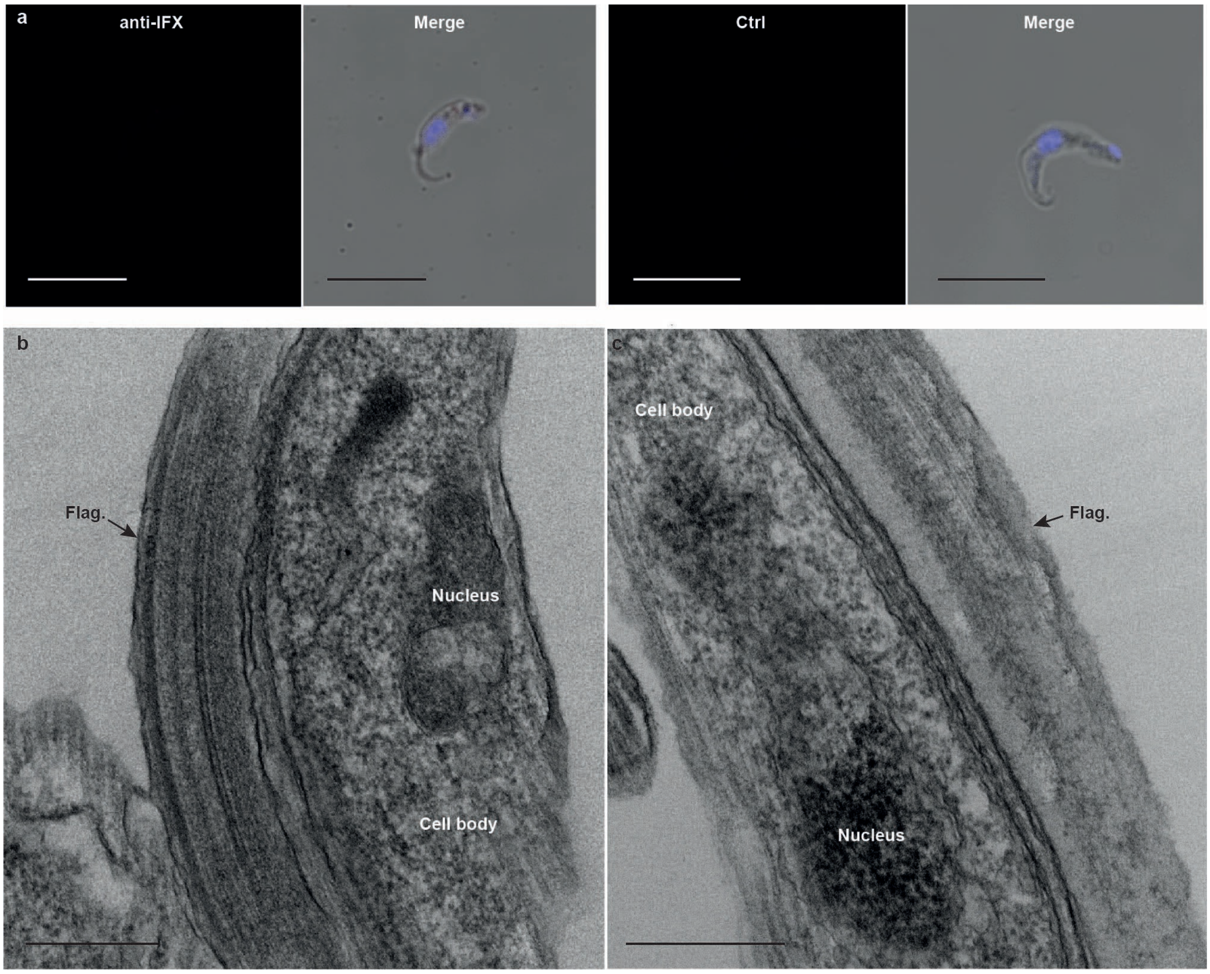


**Extended Data Figure 5 | Absence of staining in anti-IFX-exposed *T. congolense* parasites and *T. vivax* isotope-matched and secondary-only controls. a,** *T. congolense* parasites were stained with anti-IFX rabbit polyclonal sera (left) or control preimmune sera (right) followed by fluorescently conjugated anti-rabbit secondary (red) and counterstained with DAPI (blue). No staining of the parasites was observed demonstrating antibody specificity. **b,** Control electron micrographs of *T. vivax* parasites stained with an isotype-matched control mouse IgG1 antibody (left panel) or goat anti-mouse coated gold particles alone (right) showing no accumulation of gold particles. Scale bars represent 8 μm (**a**) and 150 nm (**b**). Flag. = flagellum.

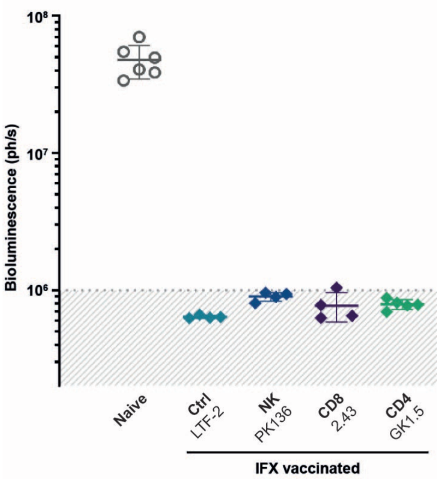


**Extended Data Figure 6 | Depletion of CD4 and CD8-positive T-lymphocytes and NK cells in IFX-vaccinated mice prior to parasite challenge do not affect IFX-mediated protective efficacy.** Groups of mice were vaccinated with IFX, rested, and then either NK cells or CD4 or CD8-positive T-lymphocytes were depleted using lineage-specific monoclonal antibodies (antibody clone names indicated) before challenging with luciferase-expressing *T. vivax* parasites. Cell-depleted animals showed no significant difference to control animals treated with an isotype-matched control antibody. The virulence of parasites was confirmed by showing robust infections in naïve mice in the same experiment. Parasitaemia was quantified on day 5 using bioluminescence, grey shading indicates bioluminescence thresholds of uninfected mice; data points represent individual animals and bars indicate mean ± SD.

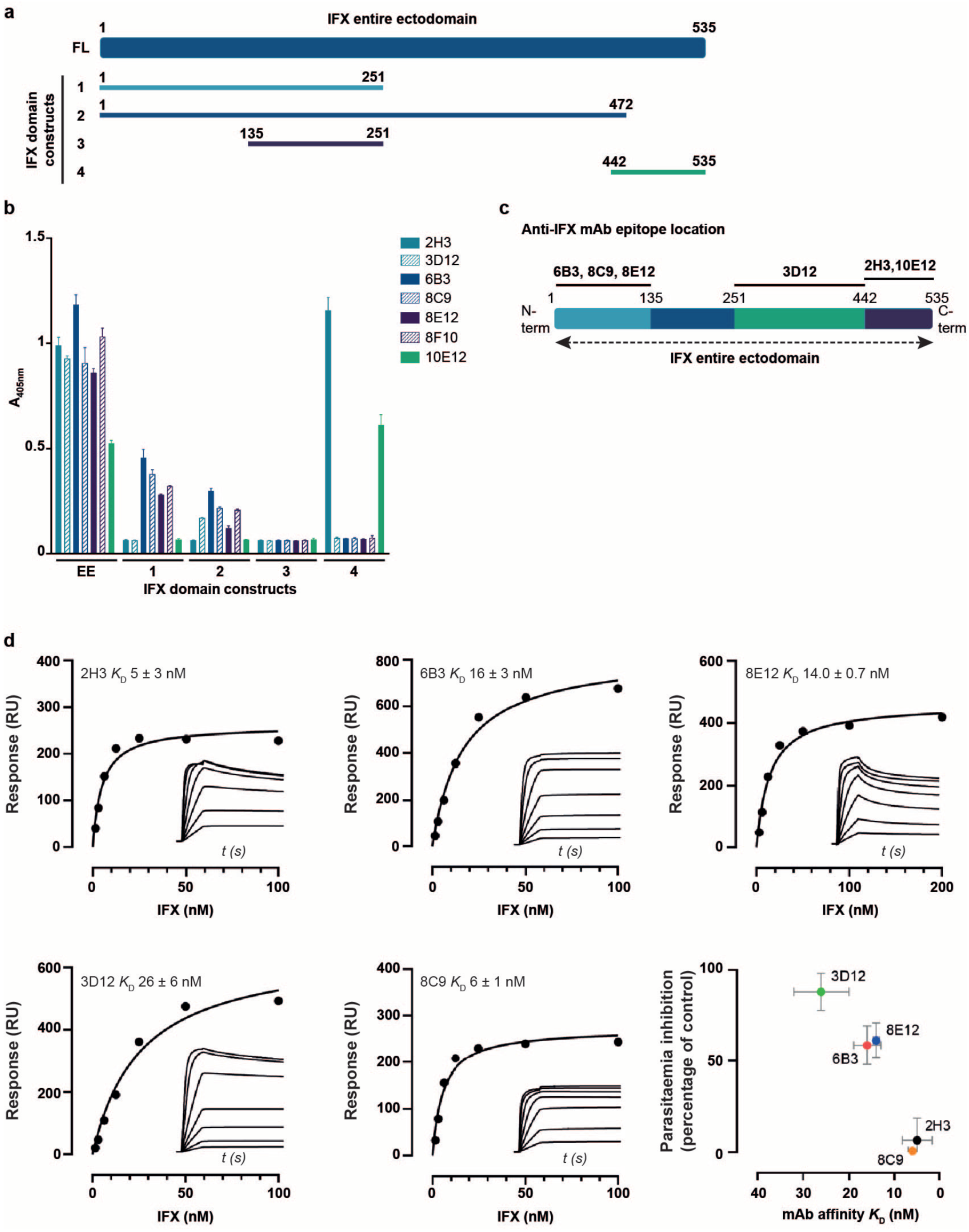


**Extended Data Figure 7 | Identification of the binding epitopes and affinities of a panel of mouse monoclonal antibodies recognising *T. vivax* IFX. a**, Schematic showing the N- and C-terminal boundaries of four fragments of the IFX ectodomain. **b**, Identification of the epitope locations for the anti-IFX monoclonal antibodies. The entire ectodomain (EE) and derived fragments (1 to 4) were expressed as enzymatically biotinylated soluble recombinant proteins in HEK293 cells, immobilised on streptavidin-coated microtitre plates, and the binding of each of the anti-IFX monoclonal antibodies quantified by ELISA. The hybridoma secreting mAb 8F10 was not successfully cloned and therefore not further investigated. Bars are mean ± SD; n ≥ 3. **c**, Schematic interpretation of the antibody binding data to the IFX ectodomain fragments showing the approximate locations of the antibody epitopes. **d**, Quantification of the equilibrium binding affinity of the anti-IFX monoclonal antibodies by surface plasmon resonance. Five of the anti-IFX monoclonal antibodies were chemically biotinylated and immobilized on a streptavidin-coated senor chip and the binding to serial dilutions of purified soluble IFX ectodomains measured. The binding affinity for each of the antibodies was calculated by fitting the binding data (inset) to a simple 1:1 binding isotherm. There was no simple positive correlation between the antibody binding affinity and protective efficacy.

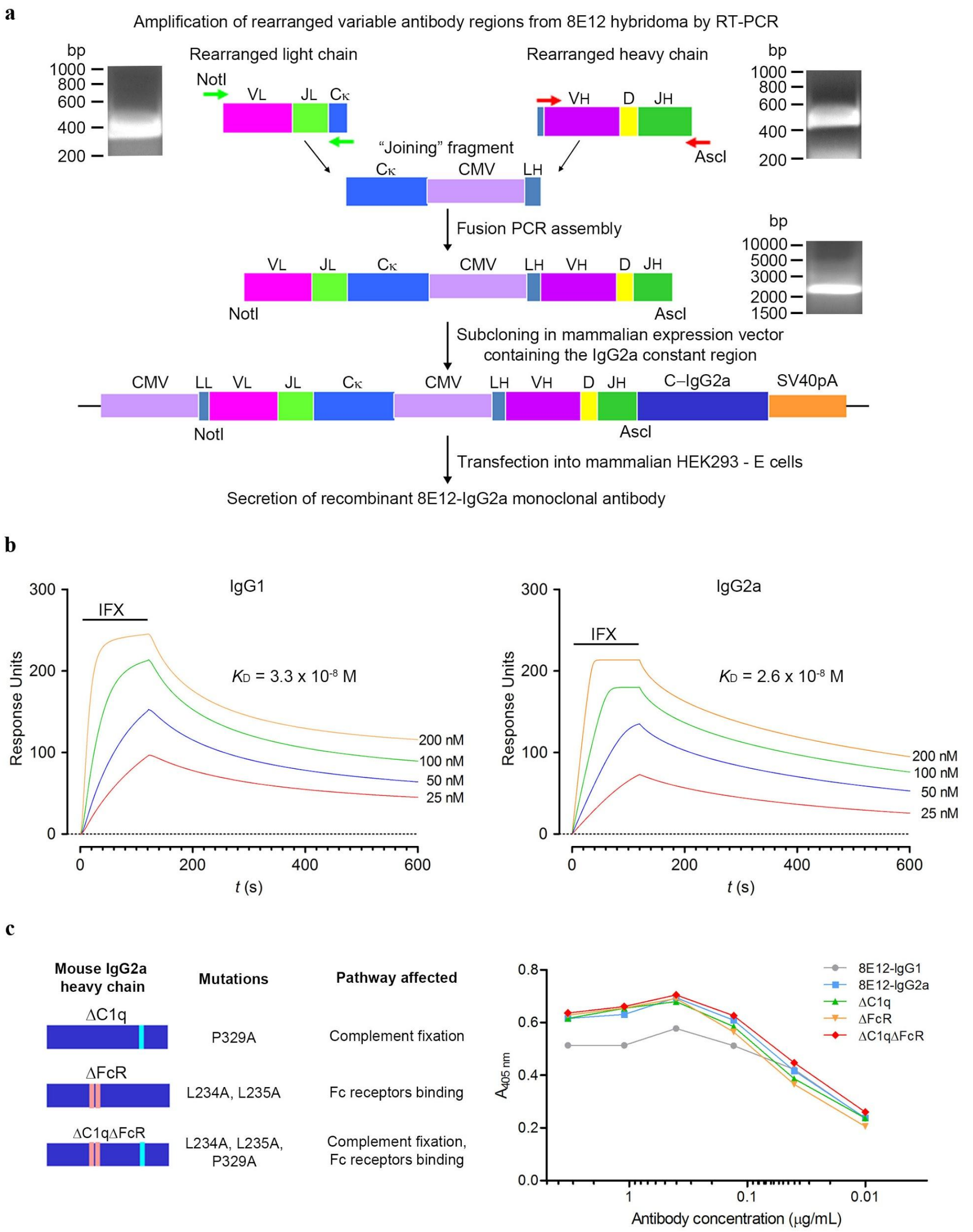


**Extended Data Figure 8 | Recombinant antibody cloning, isotype switching, and mutation of antibody effector recruitment sites of the anti-IFX 8E12 hybridoma. a**, The rearranged variable light and heavy regions of the anti-IFX 8E12 monoclonal antibody were amplified and assembled by fusion PCR using a "joining" fragment before being subcloned into a mammalian protein expression plasmid containing the mouse IgG2a heavy chain. The 8E12-IgG2a antibody was produced by transfection of HEK293 cells. **b**, The binding affinity of the 8E12 monoclonal antibody for IFX is unaffected after isotype switching. The biophysical binding parameters of the 8E12 monoclonal antibody for IFX were determined by SPR as both the hybridoma-expressed IgG1 (left panel) and recombinant IgG2a (right panel). Serial dilutions of the purified complete ectodomain of IFX were injected for two minutes over the biotinylated antibodies immobilised on a streptavidin-coated sensor chip and left to dissociate. Equilibrium binding constants were calculated by fitting the binding data to a Langmuir binding isotherm and found to be essentially equivalent. **c**, Mutation of the C1q and FcR recruitment sites on the 8E12-IgG2a heavy chain. The specified mutations which are known to abrogate binding to either C1q or FcR were made on the recombinant 8E12-IgG2a plasmid using site directed mutagenesis. Mutations were made individually (ΔC1q and ΔFcR) and together (ΔC1qΔFcR). Each of the three mutant antibodies were expressed, purified, and IFX-binding activity normalised to the parent 8E12-IgG2a and 8E12-IgG1 by ELISA.

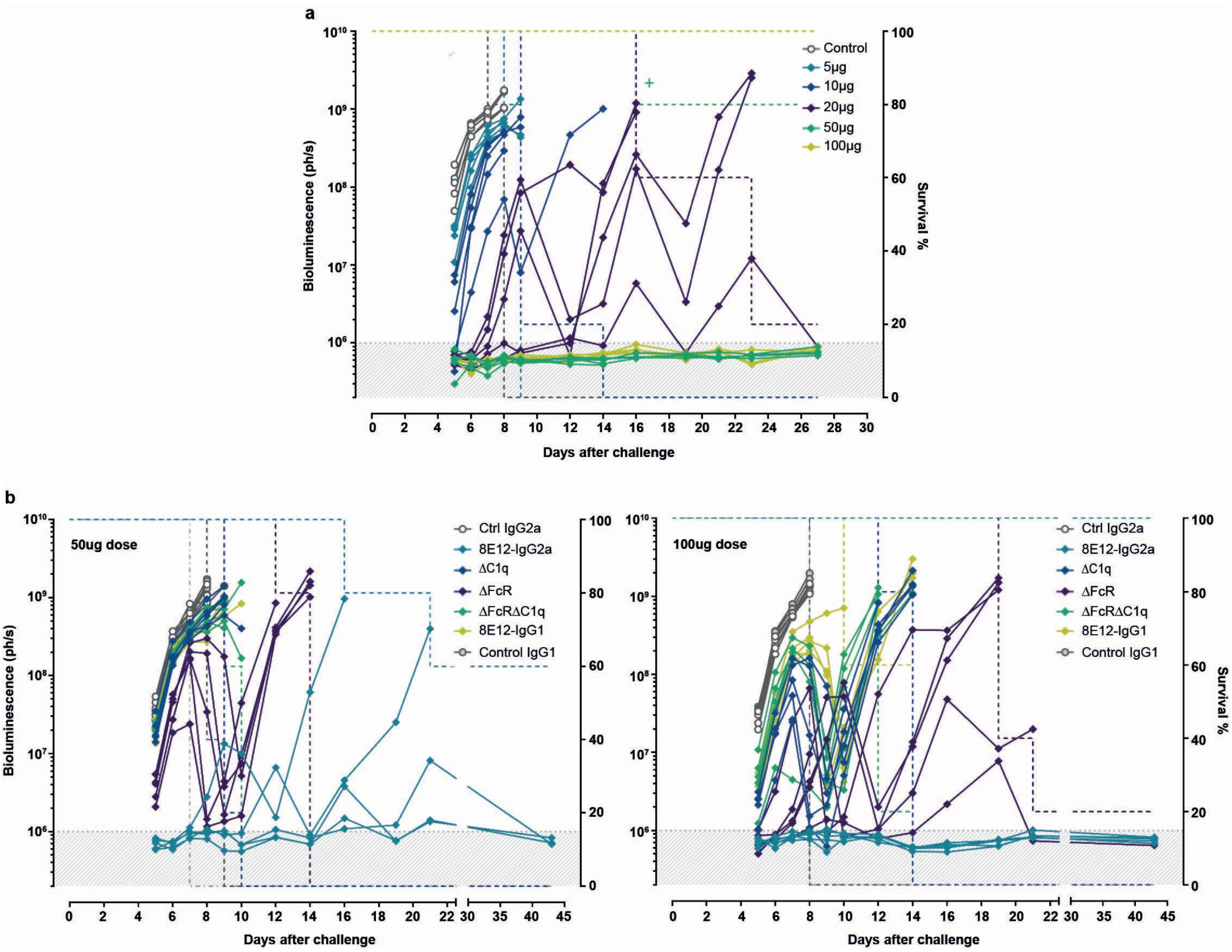


**Extended Data Figure 9 | The anti-IFX 8E12-IgG2a monoclonal antibody with abrogated immune effector recruitment sites reveals highly potent protection due to multiple mechanisms of immunological protection including a major role for complement. a,** Groups of five mice were injected three times intravenously with the indicated doses of purified anti-IFX 8E12-IgG2a monoclonal antibody and challenged with the luciferase-expressing transgenic *T. vivax* parasites. Control is an isotype-matched mouse IgG2a monoclonal antibody. A cross indicates where a single animal had to be removed from the study on day 16 for health reasons thought to be unrelated to the infection. **b,** Groups of five mice were administered three times intravenously with either 50 µg (left panel) or 100 µg (right panel) of purified anti-IFX 8E12-IgG2a monoclonal antibody containing mutations in immune effector recruitment binding sites and challenged with luciferase-expressing transgenic *T. vivax* parasites. Mutations prevented binding to C1q (ΔC1q), FcRs (ΔFcR) or both (ΔC1qΔFcR) and were compared to non-mutated 8E12-IgG2a, 8E12-IgG1 and both isotype-matched IgG2a and IgG1 controls. In all panels, data points represent individual animals and grey shading indicates bioluminescence thresholds of uninfected mice; dashed lines indicate survival within each group. Reductions in parasitaemia followed by rebounds after day 8 post infection are likely to be due to the development of protective host antibody responses directed to the dominant variable surface glycoprotein (VSG) within the parasite population and selection of an antigenically distinct variant. One of two independent experiments with very similar outcomes is shown.

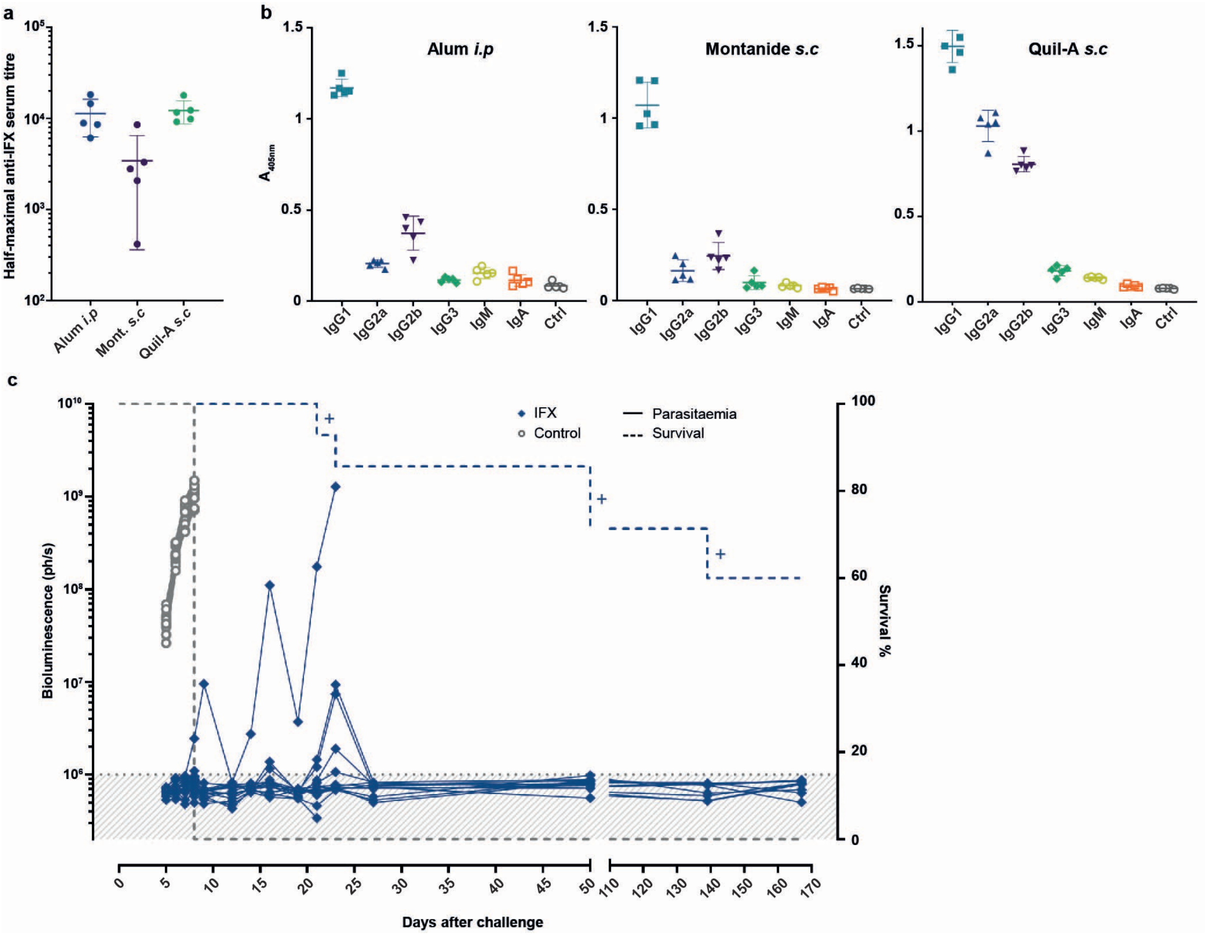


**Extended Data Figure 10 | IFX adjuvanted in Quil-A and delivered subcutaneously induces consistent and isotype-balanced anti-IFX titres that are highly protective. a,** Mice were immunised with the purified ectodomain of IFX adjuvanted in alum, Quil-A and Montanide ISA 201 VG (Mont.) using a prime and two-boost regime either intraperitoneally (alum i.p.) or subcutaneously (Quil-A and Montanide s.c.). Half-maximal anti-IFX titres were determined by ELISA. IFX/Quil-A administered subcutaneously was able to elicit anti-IFX antibody titres that were as high as IFX/alum delivered intraperitoneally. **b,** Quantification of different anti-IFX antibody isotypes elicited by the different adjuvants. IFX/Quil-A was able to induce a larger proportion of IgG2 isotype subclasses. Data points represent individual mice and bars are mean ± SD. **c,** Increased protection to *T. vivax* challenge using Quil-A in a protein-in-adjuvant vaccine formulation. Fourteen mice were immunized subcutaneously with purified soluble IFX recombinant protein adjuvanted in Quil-A and challenged with transgenic luciferase-expressing *T. vivax*. Parasitaemia was quantified on the indicated days after parasite challenge using bioluminescence; controls are a cohort of 14 animals treated with adjuvant only. Data points represent individual animals and grey shading indicates bioluminescence thresholds of uninfected mice. Crosses indicate where individuals had to be removed from the study for health reasons thought to be unrelated to the infection. Note that the smaller bioluminescence peaks in four mice corresponding to high bioluminescent readings between days 16 and 24 were caused by bleed-through of bioluminescence signal from the mouse that eventually succumbed to infection
